# Antigen receptor competition in CAR-TCR cells changes cytotoxicity outcomes upon spatial antigen separation

**DOI:** 10.1101/2025.10.07.680634

**Authors:** Carolin Moosmann, Felix Drost, Ev-Marie Schuster, Nikolaj Pagh Kristensen, Mariana Sousa, Martin Böttcher, Anna Pavlova, Vivien Reinecke, Heiko Bruns, Stefan Uderhardt, Michael Hudecek, Simon Völkl, Andreas Mackensen, Dirk H. Busch, Benjamin Schubert, Dimitrios Mougiakakos, Kilian Schober

**Affiliations:** Mikrobiologisches Institut–Klinische Mikrobiologie, Immunologie und Hygiene, Universitätsklinikum Erlangen, Friedrich-Alexander Universität (FAU) Erlangen-Nürnberg, Erlangen, Germany; Institute of AI for Health, Computational Health Center, Helmholtz Munich, Neuherberg, Germany; Helmholtz Pioneer Campus, Helmholtz Munich, Neuherberg, Germany; Department of Hematology, Oncology, and Cell Therapy, Otto-von-Guericke University, Magdeburg, Germany; Department of Internal Medicine 5 - Hematology and Oncology, FAU Erlangen-Nürnberg and Universitätsklinikum Erlangen; Erlangen, Germany; Department of Internal Medicine 3-Rheumatology and Immunology, FAU Erlangen-Nürnberg and Universitätsklinikum Erlangen, Erlangen, Germany; Deutsches Zentrum für Immuntherapie (DZI), FAU Erlangen-Nürnberg and Universitätsklinikum Erlangen, Erlangen, Germany; Exploratory Research Unit, Optical Imaging Centre Erlangen, FAU Erlangen-Nürnberg, Erlangen, Germany; Chair in Cellular Immunotherapy, Department of Medicine II, University Hospital Würzburg, Würzburg, Germany; Fraunhofer Institute for Cell Therapy and Immunology (IZI), Cellular Immunotherapy Branch Site, Würzburg, Germany; National Center for Tumor Diseases (NCT), Site Würzburg-Erlangen-Regensburg-Augsburg (WERA), and Bayerisches Zentrum für Krebsforschung (BKFZ), Site Würzburg, Würzburg, Germany; FAU Profile Center Immunomedicine, FAU Erlangen-Nürnberg, Erlangen, Germany; Institute for Medical Microbiology, Immunology, and Hygiene, School of Medicine and Health, Technical University of Munich, Munich, Germany; German Center for Infection Research, Partner Site Munich, Munich, Germany; Institute of Compuational Biology, Helmholtz Munich, Neuherberg, Germany

## Abstract

Chimeric antigen receptor T cells (CAR-T) are effective against cancer and autoimmunity, but the role of the endogenous T-cell receptor (TCR) remains unclear. Using TCRs as molecular barcodes, we traced anti-CD19 CAR-T clonotypes before and after infusion in patients with B-cell malignancies using single-cell sequencing. A cytotoxic phenotype, but not transcriptional responsiveness to tumor cells, correlated with CAR-T detection *in vivo*. To assess functional TCR contributions to CAR-T activity, we combined CAR transduction with orthotopic TCR replacement. Strong TCR signaling enhanced activation, but impaired CAR-mediated cytotoxicity when TCR and CAR antigens were presented by different target cells. Confocal imaging revealed preferential recruitment of cytolytic granules to the TCR synapse in cells simultaneously forming distinct CAR and TCR synapses. Thus, spatial antigen separation shapes receptor competition in CAR-TCR cells. Our findings have implications for therapeutic CAR-T design using defined TCRs to enhance cellular persistence or counteract tumor heterogeneity.

## Introduction

CAR-T cells are effective against cancer and autoimmunity^1,2^. Manufacturing typically introduces the CAR by viral transduction while retaining the endogenous TCR. As a molecular barcode, the TCR enables clonotypic tracking of CAR-T cells, linking cytotoxic or more differentiated infusion-product phenotypes with *in vivo* maintenance^3–6^. However, the TCR is also a functional receptor whose specificity and engagement may influence CAR-T cell behavior. Associations between phenotype and clonal maintenance may therefore reflect both the state of the tracked cells and functional contributions of their TCRs. Understanding these contributions requires studying the TCR both as a clonal marker and as a potential regulator of CAR-T function.

Evidence for TCR-mediated functional contributions comes from studies deleting the endogenous TCR. TCR deletion can prevent graft-versus-host disease caused by allogeneic CAR-T cells^7–9^, but may also remove beneficial TCR signals. TCR knock-out (KO) preserved CAR-T function *in vitro* but reduced maintenance in a xenograft mouse model, possibly reflecting a contribution of xenoreactivity to TCR^+^ CAR-T cell maintenance^7^.

Similarly, using Epstein-Barr virus (EBV) or cytomegalovirus (CMV)-specific T cells for CAR-T manufacturing has been proposed to boost CAR-T cell responses through continuous antigen exposure, and retain an effector rather than exhausted phenotype^10,11^. However, EBV-specific CAR-T cells showed diminished long-term maintenance compared to conventional polyclonal CAR-T cells within the same patients^10^. This could reflect TCR interference with CAR function, although differences in starting phenotypes and manufacturing complicate interpretation^10–12^.

A recent study directly showed that strong TCR signaling – as exerted by virus-specific TCRs – enhances CAR-T function, while weak signaling can antagonize it^13^. However, this analysis largely focused on CAR and TCR antigens presented by the same target cell. In patients, antigens may instead be presented by different cells when tumor-directed CARs are combined with virus-specific TCRs^10^, or with TCRs targeting complementary antigens in heterogeneous tumors^14,15^. Following this concept, another study showed that dual-receptor T cells preferentially killed CAR targets, when antigens were presented by separate target cells and TCR antigen levels were low^16^. Overall, in human^16^ and murine^9,17^ systems, abundant antigen for either receptor can impair killing through the other, yet it remains poorly understood how antigen presentation on the same or different target cells shapes TCR-CAR interplay.

Here, we first used TCR sequences as molecular barcodes to link anti-CD19 CAR-T phenotypes in the infusion product to clonal CAR-T detection post-infusion in patients with B-cell malignancies by single-cell sequencing (sc-seq). To separately assess functional contributions of the TCR itself on CAR-T cells, we then combined CAR transduction and CRISPR/Cas9-mediated orthotopic TCR replacement (OTR), enabling controlled evaluation of TCR presence, specificity and activity. Comparing antigen presentation by the same (*cis*) or different (*trans*) target cells revealed that strong TCR signaling impaired CAR-mediated cytotoxicity only *in trans*. This was accompanied by preferential recruitment of the cytolytic machinery to the TCR synapse during simultaneous target cell engagement.

## Results

### A subset of pre-infusion CAR-T cells shows transcriptomic shifts upon in vitro stimulation

To identify pre-infusion features associated with post-infusion CAR-T cell detection, we profiled phenotypes and clonotypes before and after tisagenlecleucel infusion in three patients with B-cell malignancies using sc-seq. Patients C14 and C20 had diffuse large B-cell lymphoma, and C16 had acute lymphoblastic leukemia (**Table S1**). To additionally assess short-term responsiveness of pre-infusion CAR-T cells at the single-cell and clonotype level, we applied “reverse phenotyping”, which identifies antigen-reactive cells through stimulation-induced transcriptomic shifts^18^: infusion-product CAR-T cells were stimulated with CD19-expressing Nalm6 cells prior to sc-seq, and compared with unstimulated cells (**Fig. 1A**).

**Figure 1:**
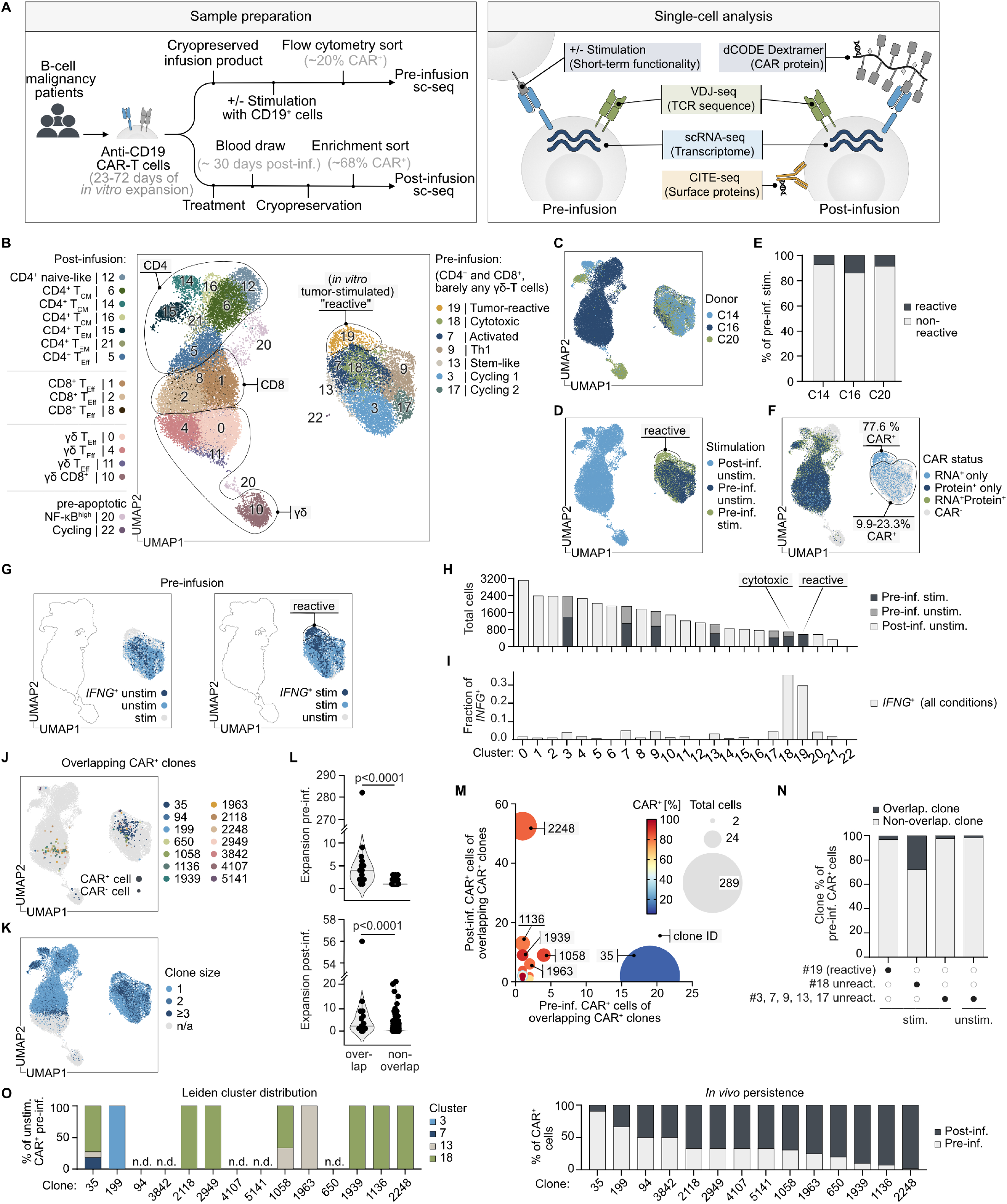
A cytotoxic pre-infusion phenotype and not short-term responsiveness to tumor cells correlates with post-infusion CAR-T cell detection. **(A)** Experimental setup (left) and single-cell analysis (right). **(B)** Leiden clusters. **(C)** Donor distribution. **(D)** Stimulation conditions. **(E)** Quantification of stimulated pre-infusion cells per donor inside and outside the reactive cluster 19. **(F)** CAR status based on CAR gene expression (RNA^+^) and/or detection of dCODE-Dextramers (Protein^+^; only in post-infusion samples). Percentages indicate CAR RNA^+^ in pre-infusion cells (reactive cluster vs. other clusters). **(G)** *IFNG* detection in unstimulated (left) and stimulated (right) pre-infusion cells. **(H-I)** Absolute cell numbers (H) and fraction of *IFNG*^+^ cells (I) per Leiden cluster. **(J)** Overlapping CAR^+^ clonotypes (cells with the same TCR sequence with ≥1 CAR^+^ cell pre- and post-infusion). **(K)** Clone sizes. **(L)** Pre-infusion (top) and post-infusion (bottom) cell numbers of overlapping and non-overlapping clonotypes. Statistical testing by Mann-Whitney-U-Test. **(M)** Absolute numbers of CAR^+^ cells of overlapping CAR^+^ clones pre- and post-infusion. **(N)** Proportions of pre-infusion CAR^+^ cells belonging to overlapping or non-overlapping clonotypes within the indicated Leiden clusters and stimulation conditions. **(O)** Leiden cluster distribution of unstimulated pre-infusion CAR^+^ cells of overlapping CAR^+^ clones (left) and proportion of pre- and post-infusion CAR^+^ cells out of all CAR^+^ cells within these clones (right); n.d., not detected.

Infusion-product CAR-T cells were identified by CAR RNA expression. Post-infusion samples were collected on day 26, 41 or 44 after CAR-T treatment and enriched for CAR^+^ cells by sorting (**Fig. 1A**). Post-infusion samples were additionally stained with CITE-seq antibodies targeting CD4, CD8 and other phenotypic markers, including DNA-barcoded CD19 CAR Dextramers to detect CAR protein surface expression (**Fig. S1A**).

Pre- and post-infusion cells occupied distinct Leiden clusters (**Fig. 1B, Table S2**) with CD4^+^, CD8^+^ and double-negative cells (**Fig. 1C; Fig. S1A-B**). Pre-infusion cluster 19 emerged upon Nalm6 stimulation (**Fig. 1D-E; Fig. S1C-D**) and was enriched for CAR RNA^+^ cells (**Fig. 1F**). Post-infusion, CAR RNA was detectable only in cells with high surface CAR protein expression (**Fig. S1E**). Consistent with previous reports^5,6,19^, post-infusion CAR-T cells were enriched for CD4/CD8-double-negative γδ-T cells (**Fig. S1F-G**). Among CAR protein-positive cells, γδ-T cells less frequently had detectable CAR RNA than αβ-T cells (**Fig. S1H**).

The reactive cluster 19 was enriched for *IFNG*-expressing cells following stimulation (**Fig. 1G-I**). Cluster 18 also expressed *IFNG*, but contained more similar proportions of stimulated and unstimulated cells (**Fig. 1H-I**). Differential gene expression analysis distinguished a cytotoxic program in cluster 18 (*CCL5, GNLY* and *GZMB*) from an activation-associated program in cluster 19 (*IL2RA* and NF-κB-related genes; **Table S2**). Activated cluster 7 and cycling clusters 3 and 17 likewise contained similar proportions of stimulated and unstimulated cells, supporting the cluster 19 phenotype as the specific transcriptional response to CD19 stimulation.

### A cytotoxic pre-infusion phenotype with limited transcriptional responsiveness correlates with post-infusion CAR-T cell detection

We next asked whether transcriptional responsiveness to tumor cells predicted post-infusion CAR-T detection. Fourteen clonotypes contained CAR^+^ cells both before and after infusion (**Fig. 1J**). These overlapping clones co-clustered with expanded clones and were larger than non-overlapping clones, particularly pre-infusion (**Fig. 1K-L**), raising the possibility that larger clones were simply more likely to be sampled twice. Yet, comparing CAR^+^ cell numbers within overlapping clones revealed distinct trajectories (**Fig. 1M**): the large clone 35 contained relatively few CAR^+^ cells and was not enriched post-infusion, whereas the smaller clone 2248 showed disproportionate post-infusion enrichment with many CAR^+^ cells. These observations suggest differential clonal dynamics beyond initial clone size, although sampling effects cannot be excluded.

We then asked whether recruitment into the reactive cluster 19 after stimulation predicted post-infusion detection. Unexpectedly, pre-infusion CAR^+^ T cells belonging to overlapping clones were enriched in the cytotoxic cluster 18 rather than the reactive cluster 19 (**Fig. 1N**). Focusing on unstimulated infusion-product cells, we again found an enrichment for the cluster 18 phenotype among overlapping clones, irrespective of their initial size or relative representation before and after infusion (**Fig. 1O; Fig. S1I**).

In summary, harnessing the endogenous TCR for CAR-T clone tracking confirmed previous findings that a cytotoxic phenotype in the infusion product was associated with *in vivo* persistence. Intriguingly, this favorable cell program was characterized by a certain inertness to *in vitro* stimulation with tumor cells, while CAR-T clones that responded strongly to tumor cells *in vitro* were no longer detectable after infusion.

### A CAR-TCR platform enables investigation of the functional impact of defined TCRs on CAR-T cells

Beyond clone size and phenotype, CAR-T cell fate may also depend on the presence, specificity, and activity of the endogenous TCR. However, isolating these relationships is challenging since natural T-cell clones differ phenotypically. To introduce defined TCRs under physiological regulation, we combined CAR transduction with CRISPR/Cas9–mediated OTR^20,21^ in primary human T cells (**Fig. S2A; methods**).

We employed this CAR-TCR platform to re-express patient-derived TCRs isolated before and after CAR-T infusion to generate sort-purified CAR-TCR cells (**Fig. S2B**). For comparison, we generated CAR^+^ TCR-KO cells (CAR-only), and CAR-T cells retaining polyclonal endogenous TCRs (CAR-endoTCR). Co-culture with CD19-expressing Nalm6 target cells induced activation and cytotoxicity in all effector populations (**Fig. S2C**), consistent with previous data^7,22^. Thus, TCR presence or identity did not affect CAR-T function under these conditions.

Because the patient-derived TCR antigen specificities were unknown, we could not use them to study how TCR engagement affects CAR-T function. We therefore engineered effector cells with a TCR recognizing the CMV epitope A*02/NLV. Purity-sorted CAR-only, CAR-TCR or TCR-only cells were co-cultured with CD19^+^ Nalm6 cells and peptide-pulsed HLA-A*02^+^ K562 cells presenting the TCR epitope (**Fig. 2A, Fig. S2D**). Apart from the NLV wild-type (WT) epitope, we tested altered peptide ligands (APLs; Mut1 and Mut2), that induce progressively weaker TCR activation (**Fig. S2E**)^23^.

**Figure 2:**
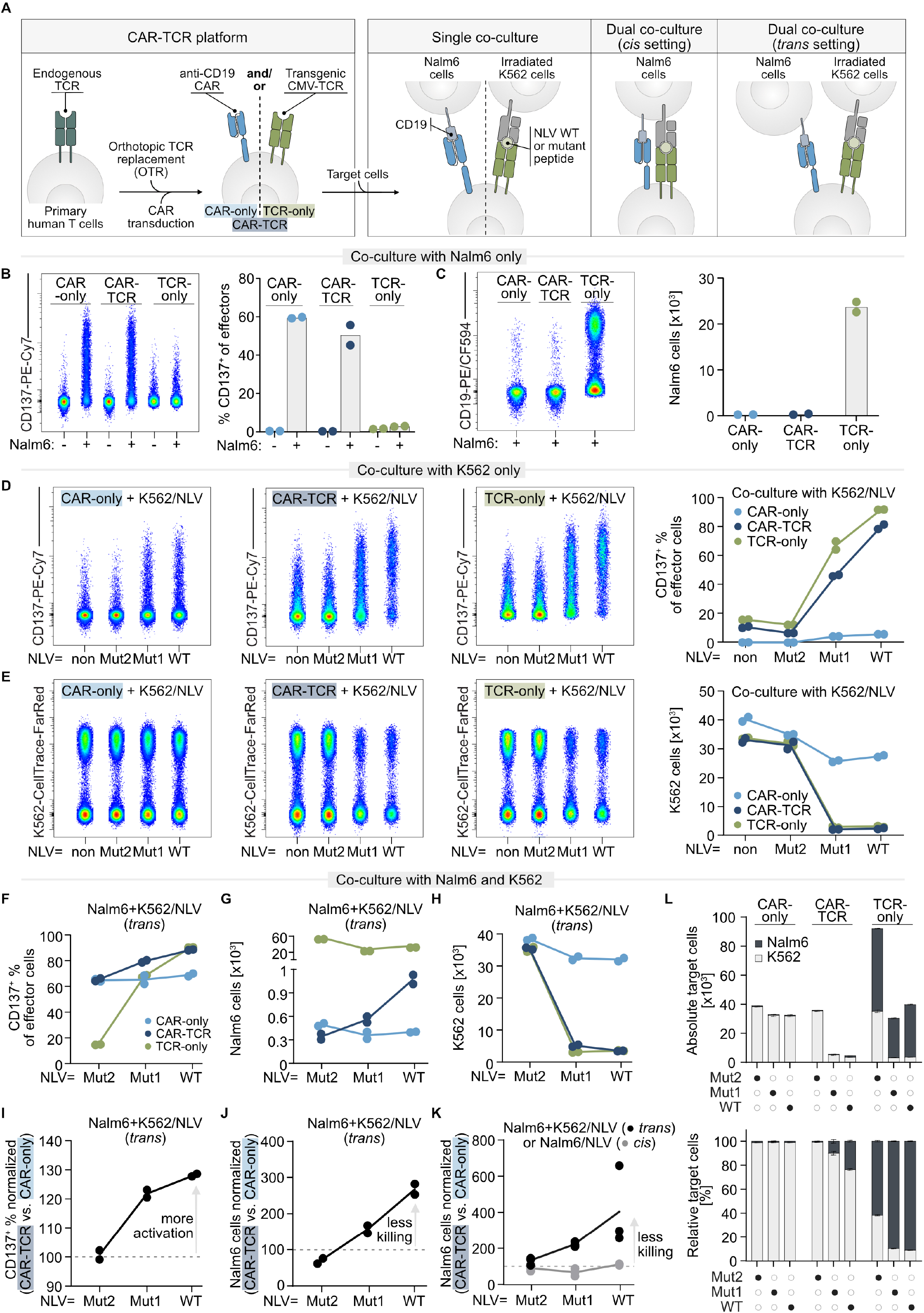
Strong TCR activity in CAR-TCR cells impairs CAR cytotoxicity against separate target cells. **(A)** Replacement of the endogenous TCR (via OTR) with an A*02/NLV CMV-specific transgenic TCR combined with anti-CD19 CAR transduction. Co-culture setups with Nalm6 (+/-NLV peptide) and peptide-pulsed K562 target cells. Effector-to-target (E:T) ratios are 2:1:1 (*trans*) or 2:1 (*cis*). **(B-E)** Flow cytometric analysis of CD137 expression in effector cells (B,D) and absolute Nalm6 (C) and K562 numbers (E) after 24 h co-culture of effector cells with only Nalm6 cells (B,C) or only peptide-pulsed K562 cells (D,E). **(F-H)** CD137 expression in effector cells (F) and absolute numbers of Nalm6 (G) or peptide-pulsed K562 cells (H) after 24 h co-culture of effector cells with both target cells (*trans*). **(I-J)** CAR-TCR vs. CAR-only ratio of CD137^+^ effector cells (I) or absolute numbers of Nalm6 cells (J) after *trans* co-cultures as in (F-H). **(K)** CAR-TCR vs. CAR-only ratio of absolute Nalm6 numbers following co-cultures with peptide-pulsed K562 and Nalm6 cells (*trans*) or peptide-pulsed Nalm6 cells only (*cis*). Statistical testing by Šídák’s multiple comparisons test (WT *trans* vs WT *cis*: 0.002, *trans* Mut2 vs *trans* WT: 0.01, remaining comparisons: ns). **(L)** Absolute (top) and relative (bottom) K562 and Nalm6 numbers after 24 h *trans* co-culture with effector cells. Data are representative of two independent experiments (B-L) and data points represent technical replicates (n=2 (B-J,L) or n=3 (K)). Bars represent mean +/-s.d. (B-C,L) and lines represent means (D-K).

We first tested antigen receptor function with each target population separately. Nalm6 cells activated CAR-expressing cells (**Fig. 2B**). CAR-only and CAR-TCR cells showed comparable killing, indicating that TCR presence alone did not impair CAR-mediated cytotoxicity (**Fig. 2C**). Conversely, NLV epitope-pulsed K562 cells activated TCR-expressing cells (**Fig. 2D**). TCR-driven cytotoxicity remained unaffected in the presence of the CAR (**Fig. 2E**). The WT NLV epitope and the APLs Mut1 and Mut2 induced progressively weaker T-cell activation (**Fig. 2D**), but cytotoxicity followed a more binary pattern: WT and Mut1 triggered similar killing, whereas Mut2 did not (**Fig. 2E**). Overall, we established a CAR-TCR platform that allows us to study the functional and molecular impact of defined TCRs on CAR-T cells.

### Strong TCR activity impairs CAR-cytotoxicity when antigens are presented by distinct cells

Next, we tested effector cells against both target populations simultaneously (*trans* setting): Nalm6 cells expressing the CAR target and K562 cells presenting the TCR target. CAR-only cells retained activation and Nalm6 cytotoxicity (**Fig. 2F-G**). TCR-only cells exhibited increasing activation depending on the TCR epitope and killed K562 cells presenting WT NLV or Mut1 (**Fig. 2F,H**).

Co-stimulation through both receptors increased CAR-TCR cell activation compared to CAR-only cells (**Fig. 2F,I**), echoing findings from a murine model where CAR and TCR targets were presented on the same cell^13^. TCR-mediated killing of K562 cells remained unaffected by CAR signaling (CAR-TCR versus TCR-only; **Fig. 2H**). Remarkably, however, increasing TCR activation strength progressively reduced CAR-mediated Nalm6 killing (CAR-TCR versus CAR-only; **Fig. 2G,J**).

We asked whether TCR engagement impaired CAR-mediated killing specifically when target antigens were presented by different cells (*trans*), which could explain differences from a recent study that focused on same-cell *cis* presentation^13^. Indeed, direct comparison of CAR-TCR and CAR-only cells revealed impaired Nalm6 killing only *in trans* (**Fig. 2K**). This impairment was already evident with the less stimulatory Mut1 epitope and was reproduced at a lower effector-to-target ratio (**Fig. S2F-I**).

We then examined the consequences of this receptor interference for therapeutic strategies targeting heterogeneous tumor cell populations (expressing only one antigen per cell type). Comparing absolute and relative numbers of surviving target cells showed that CAR-TCR cells controlled the combined heterogeneous target population better than single-receptor effectors (**Fig. 2L**). However, increasing TCR stimulation led to higher relative Nalm6 survival, reflecting worse CAR-mediated cytotoxcity.

Together, dual-receptor effector cells improved control of heterogeneous target populations, but stronger TCR stimulation compromised CAR-mediated killing.

### Strong TCR signaling diverts the cytolytic machinery from CAR immunological synapses

TCR activity could impair CAR-mediated killing by diverting effector cells and/or molecules away from CAR targets^13^. We investigated this possibility in a short-term co-culture assay followed by flow cytometric detection of effector-target cell doublets. CAR-TCR and CAR-only cells bound Nalm6 cells at similar frequencies in both *cis* and *trans* settings. Despite some non-specific binding of CAR-only and TCR-only cells to K562 and Nalm6 cells, respectively, a substantial fraction of effector cells simultaneously bound both target types, forming effector:target:target triplets (**Fig. S3A**,**B**). Thus, reduced CAR-mediated cytotoxicity was not accompanied by reduced cellular binding to CAR targets.

This prompted us to analyze intracellular interference during simultaneous receptor engagement. We examined immunological synapse organization by confocal microscopy after co-culturing CAR-TCR cells with Nalm6 and peptide-pulsed K562 cells (*trans*). We visualized actin, CARs, TCRs and the cytolytic granule component perforin. CAR-TCR cells formed synapses with either target individually, characterized by enrichment of the corresponding receptor and actin at the contact site (**Fig. 3A**). Both synapse types showed comparable perforin recruitment when confronted only with their cognate target (**Fig. 3A; Fig. S3C**).

**Figure 3:**
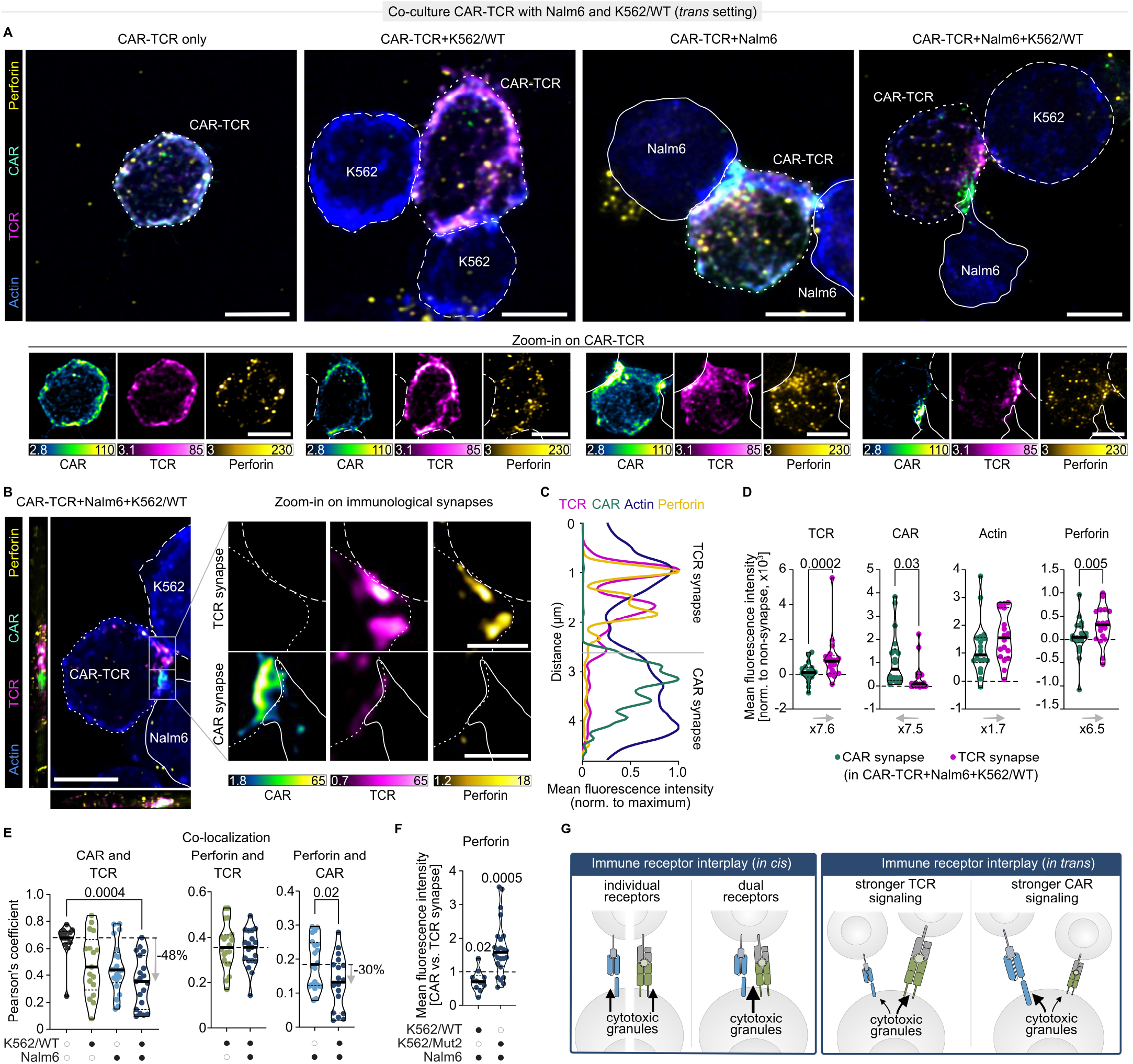
Strong TCR signaling diverts the cytolytic machinery away from CAR immunological synapses. **(A)** Confocal microscopy images of CAR-TCR cells interacting with target cells in short-term (30 min) *trans* co-culture. Solid white line: Nalm6; dashed line: K562 carrying WT NLV peptide (K562/WT); dotted line: CAR-TCR cells. Images show sum intensity projections in the z dimension. Heatmap values (lower panel): gray value ×10^3^. **(B)** Confocal microscopy images, visualized as in (A). Left, overview sum intensity projections in x, y, z planes. Right, zoom-ins of TCR (top) and CAR (bottom) synapses represent individual z-slices of the region indicated by the box on the left. Heatmap values: gray value ×10^3^. **(C)** Signal intensities within the synapse regions indicated by the box in (B). Values were normalized to the maximum for each marker. **(D)** Marker signal intensities in synapses of CAR-TCR cells interacting with both target cells simultaneously. Values were normalized to non-synaptic regions by subtraction. Statistical testing by paired t test. Percentages at the bottom indicate the signal loss/gain between medians of the two synapses, with the start of the arrow indicating the reference. **(E)** Co-localization analysis of synapse components within whole CAR-TCR cells from images as in (A). Statistical testing by Kruskal-Wallis-test (left) or unpaired t test (middle, right). Percentages on the right indicate the signal loss between medians of effector-only (left) or effector:Nalm6 conjugates (right) and heterotypic triplets. **(F)** Ratios of perforin signals in CAR vs. TCR synapses of the same CAR-TCR cell interacting with Nalm6 cells and WT or Mut2 peptide-pulsed K562 cells analyzed as in (A). Statistical testing by One sample t test (versus 1). **(G)** Graphical summary. Scale bars: 5 µm (A, B left), 1.5 µm (B right). Data are representative of one (F) or two independent experiments (A-E) and data points represent individual cell conjugates (n=18 (triplets in D-E), n=13 (effector-only in E), n=17 (effector:K562 in D-E), n=20 (effector:Nalm6 in D-E), n=10 (K562/WT in F), n=24 (K562/Mut2 in F)).

Next to homotypic synapses (i.e., one effector and one target type), CAR-TCR cells also engaged both target types simultaneously, forming distinct heterotypic synapses enriched for TCR at the K562 interface and CAR at the Nalm6 interface. Despite simultaneous formation of both synapses, perforin preferentially localized to the TCR synapse (**Fig. 3A-D; Fig. S3C**). Whole-cell colocalization analysis supported this finding: CAR–perforin colocalization was lower in heterotypic triplets than in cells engaging Nalm6 cells alone, whereas TCR–perforin colocalization was unaffected by simultaneous CAR-target engagement (**Fig. 3E; Fig. S3D**).

To assess the role of TCR stimulation strength, we repeated the imaging with Mut2-pulsed K562 cells. CAR-TCR cells still formed distinct CAR and TCR synapses, but perforin recruitment to the CAR synapse remained unaffected (**Fig. S3E-F**) and perforin levels were now higher at the CAR than at the TCR synapse (**Fig. 3F**).

Taken together, our data suggest that the “stronger” immune receptor interferes with cytotoxicity mediated by the “weaker” receptor by diverting the cytolytic machinery to its own immunological synapse, specifically when effector cells form heterotypic synapses with different target cells simultaneously (**Fig. 3G**).

## Discussion

Here, we first used the TCR as a molecular barcode to track CAR-T clones in cancer patients. We then investigated whether the TCR intrinsically influences CAR-T function in the absence or presence of signaling, and in settings of *cis* or *trans* antigen presentation.

Consistent with previous reports^3,4,6^, a cytotoxic infusion-product phenotype was associated with subsequent CAR-T cell detection *in vivo*. However, it has to be noted that the infusion product is synthetic, with T cells having undergone weeks of rapid expansion *in vitro*. Physiological immune memory formation is mainly driven by quiescent stem-like T cells^24^. Accordingly, stem-like cells in the leukapheresis product directly *ex vivo*, but not in the later CAR-T infusion product, correlate with *in vivo* persistence and clinical responses^25–27^. More strikingly, we observed that transcriptional responsiveness to tumor cell stimulation was not associated with post-infusion detection. Highly responsive cells might undergo activation-induced cell death *in vivo*, whereas less reactive cells survive better. These results need to be confirmed in larger and more homogeneous patient cohorts. Furthermore, future work needs to assess whether the *in vivo* enrichment of γδ-T cells results from functional advantage or selective maintenance, although the enhanced survival of patients with high γδ-CAR T-cell counts^6^ suggests intrinsic functional benefits.

We here established a CAR-TCR platform to systematically investigate CAR-TCR cross-talk upon *cis* or *trans* antigen presentation. Previous studies showed that weak TCR signaling antagonizes CAR-mediated killing whereas strong signaling enhances it when both antigens are presented by the same cell^13,17^. In our *cis* setting, CAR-TCR and CAR-only cells showed comparable killing. *In trans*, however, increasing TCR-engagement compromised CAR-mediated cytotoxicity. Conversely, CAR engagement did not impair TCR-mediated killing. The reduction in CAR-mediated killing was not accompanied by reduced binding to CAR targets. Instead, perforin preferentially localized to the TCR synapse in heterotypic triplets under strong, but not weak, TCR stimulation. These findings suggest that relative receptor stimulation strength influences the allocation of the cytolytic machinery between competing synapses.

It has long been known that T cells can form immunological synapses with two target cells simultaneously and cytotoxicity is not restricted to strong interactions^28,29^. However, such previous studies investigated only TCR-mediated synapses whereas we here studied CAR-TCR cells forming heterotypic synapses. We observed that stimulatory TCR synapses can outcompete CAR synapses. This may be explained by CAR synapses forming faster^24,30^ whereas TCR signaling being more sustained^17,30^. However, CAR engagement can also interfere with TCR cytotoxicity in other settings^16^. These seemingly contradicting findings are reconciled by the fact that strong but not weak TCR stimulation impaired CAR-mediated cytotoxicity. This is consistent with the concept that relative synapse strength, rather than receptor identity alone, determines preferential recruitment of cytolytic granules. Our imaging provides a spatial basis for this model of receptor competition.

Our work is relevant for therapeutic contexts in which spatial target cell separation is likely. If target cells are far away from each other, we do not expect synapse competition (e.g., CAR target and TCR target present in different organs). Conversely, if antigens are present on the same cell (*cis* setting), cooperative signaling through CARs and TCRs might be expected. However, when CAR and TCR antigens are present on neighboring cells, immune receptor competition can occur. Notably, this may be particularly relevant for tumors with heterogeneous antigen expression, which represent the very setting for which complementary tumor-specific TCRs and CARs on the same effector cell are intended^14,15^. CAR-TCR interference may further occur if virus-specific TCRs are combined with tumor-specific CARs, and virus-positive cells are present next to tumor cells. For clinical settings like the one by Li et al., in which anti-GD2 CAR-T cells with EBV-specific TCRs showed reduced long-term maintenance^10^, the exact topology of target cells may be a critical determinant. Our findings suggest that spatial target cell distribution should be considered in CAR-TCR therapy design.

## Methods

### Study design

The aim of the study was to (1) identify factors influencing CAR-T persistence in cancer patients and (2) determine whether an endogenous-like TCR and a transduced CAR interfere in CAR T cells. We used samples from three cancer patients and additionally performed controlled laboratory experiments on genetically engineered T cells from healthy donors. Data representing controlled laboratory experiments were not subjected to sample blinding but compared cell populations were generated from the same donor T cells and treatments were run in parallel.

### Study cohort

Ethics approvals (number 219_14B, number 336_19B) were granted by the local Ethics Committee of the Medical Faculty of the University Hospital of Erlangen, Friedrich-Alexander Universität Erlangen-Nürnberg, Germany. Samples were collected after informed consent of the donors. Three female donors (C14, C16 and C20) aged 62, 19 and 69, respectively, received anti-CD19 CAR-T cells (Kymriah, Novartis). For the pre-infusion product, leftover cells were rinsed from the infusion bag and tubes. Peripheral blood mononuclear cells (PBMCs) were isolated from citrated peripheral blood by density gradient centrifugation using a BioColl density medium with a density of 1.077 g/mL (BioSell, BS.L 6115). Cells were resuspended in heat-inactivated FCS + 10% DMSO and stored in liquid nitrogen. For the post-infusion samples, blood was drawn on day 41 (C14), day 26 (C16) or day 44 (C20) and PBMCs were isolated and cryopreserved.

### Stimulation of pre-infusion CAR-T cells for reverse phenotyping

Thawed pre-infusion CAR-T cells and Nalm6 cells were washed in medium (in RPMI 1640 + 10% FCS + 2 mM L-glutamine + 25 mM HEPES + 20 IU/mL Interleukin-2 (IL-2)). After manual counting, 2 mL CAR-T cells of each donor (5×10^5^ cells/mL) were either co-cultured with 0.5 mL Nalm6 (1×10^6^ cells/mL) (=stimulated) or 0.5 mL of additional medium (=unstimulated), respectively, in 6-well plates at 37°C, 5% CO_2_ for 24 h.

### Single-cell RNA sequencing (scRNAseq) of pre-infusion CAR-T cells

Stimulated and unstimulated cells were subjected to flow-based cell sorting. After an initial washing step with PBS (300 xg for 6 min at 21°C), samples were incubated with ZombieAqua (1:500, BioLegend) for 10 min in the dark at room temperature. Samples were washed once with 1 mL PBA (PBS + 0.25% BSA + 0.02% NaN_3_) and human CD19 CAR Detection Reagent (1:50, Miltenyi biotec, 130-129-550) was added, mixed and incubated for 10 min at room temperature in the dark. After washing twice with 2 mL PBA at room temperature, an antibody mastermix comprising anti-CD19-PE antibody (1:50, clone 4G7, BioLegend), anti-CD3-APC (1:33, UCHT1, BioLegend) and anti-Streptavidin-PE/TexasRed (1:200, ThermoFisher, SA1017) was added and incubated for 20 min at room temperature in the dark, followed by two final washing steps in sorting buffer (PBA + 1 mM EDTA). Single, live CD19^-^ CD3^+^ lymphocytes (8333 cells per donor and condition) were sorted on a MoFlo XDP (Beckman Coulter) sorter. Immediately after sorting, cells were pooled into one 10X sample according to their stimulation condition and loaded to a Chromium Next GEM Chip K (10X Genomics) and Chromium Next GEM Single-Cell 5′ kits v.2 were used to generate gene expression (GEX) and T cell receptor (VDJ) libraries according to the manufacturer’s instructions (10X Genomics, 1000266, 1000196, 1000256, 1000252, 1000287, 3000431, 3000510). Libraries were sent to Novogene and sequenced on an Illumina NovaSeq platform with PE150 strategy.

### scRNAseq of post-infusion CAR-T cells

Cryopreserved post-infusion CAR-T cells were thawed and rested overnight at 2×10^6^ cells/mL in cRPMI (RPMI 1640 Medium + 10% heat-inactivated FCS, 0.05 mM β-mercaptoethanol, 0.05 mg/mL gentamicin, 1.1915 g/L HEPES, 0.2 g/L L-glutamine, 100 U/mL Penicillin-Streptomycin). After manual counting, all cells (1.35×10^6^ for C14, 3.55×10^6^ for C16, 3.1×10^6^ for C20 cells) were collected and washed twice in 200 µL sterile-filtrated Wash buffer (PBS, pH 7.4, with 1% filtered FCS) and then resuspended in the 50 µL (C14) or 100 µL (C16, C20) sterile-filtrated Stain buffer (PBS, pH 7.4 containing 1% FCS, 0.1 g/L Herring sperm DNA). 4.4 µL (C14) or 8.8 µL (C16, C20) of dCODE reagent mastermix (2.0 or 4.0 μL of CD19-dCODE® Dextramer (Immudex, DS005), 2.0 or 4.0 μL of control-dCODE® Dextramer (Immudex, WI03233), and 0.4 or 0.8 μL 100 μM d-Biotin) were added to each sample. After 15 min incubation in the dark at room temperature, mastermixes comprising a DNA-barcoded hashtag antibody (0.5 µL per 1×10^6^ PBMCs of TotalSeq-C anti-human hashtag antibodies 1–3 (BioLegend 394661, 394663, 394665), DNA-barcoded CITE-Seq TotalSeq-C antibodies (per 5×10^6^ PBMCs: 0.25 µg of anti-human CD4 (BioLegend, 300567), 0.25 µg anti-human CD8 (BioLegend, 344753), 0.078 µg anti-human CD45RA (BioLegend, 304163), 0.078 µg anti-human CD62L (BioLegend, 304851), 0.3125 µg anti-human CD95 (BioLegend, 305651), 0.277 µg anti-human CCR7 (BioLegend, 353251) in Stain buffer (PBS, pH 7.4, with 1% FCS and 0.1 g/L Herring sperm DNA) and surface antibodies and viability dye (anti-CD56-FITC (1:200, Invitrogen, 11-0566-42), anti-CD19-FITC (1:200, BioLegend, 302205), anti-CD4-BV711 (1:400, BD Horizon, 568371), anti-CD3-PE-Cy7 (1:200, Beckman Coulter, 737657), anti-CD8a-APC (1:400, BioLegend, 301014), anti-CD14-APC/Cy7 (1:200, BD Pharmingen, 557831) and ZombieAqua (1:500, BioLegend, 423102) in Stain buffer) were centrifuged (14,000 xg, 10 min, room temperature) and then added to the respective cell suspension. After 20 min incubation in the dark on ice, samples were washed 4 times with Wash buffer (300 xg, 5 min, 4°C) and then resuspended in 500 µL sterile-filtered FACS buffer (PBS, 0.5% BSA). Sort enrichment was performed for single, live, CD56^-^, CD19^-^, CD14^-^, CD3^+^ dCODE-Dextramer^+^ cells on a MoFlo Astrios EQ sorter (Beckman). In total, 5,670 C14 cells, 69,925 C16 cells and 11,560 C20 cells were sorted into pre-coated 1.5 mL tubes containing FACS-buffer. Immediately after sorting, scRNAseq samples were prepared and sequenced as described above. In addition to GEX and VDJ libraries, a cell surface (CITEseq) library was prepared (10X Genomics, 1000266, 1000196, 1000256, 1000252, 1000287, 3000431, 3000510).

### Computational single-cell RNA sequencing data analysis

The CD19-BBz CAR gene sequence obtained from patent EP3214091 (**Table S3**) was added to the GRCh38-2020-A genome via the cellranger (10x Genomics, version 6.0.2) ‘mkref’ command. The resulting reference genome, the VDJ reference vdj-GRCh38-5.0.0, and custom feature barcodes (post-infusion samples only) were used to process the individual sequencing runs with the cellranger ‘multi’-command. Single-cell analysis was performed as described by Heumos et al.^30^ via Scanpy-1.8.2^31^ and Scirpy-0.10.1^32^.

Quality Control was conducted separately for pre- and post-infusion data as follows. All gene expression counts and TCR contig annotations were joined across the sequencing runs. Dying cells and doublets were filtered by thresholds on high fractions of mitochondrial counts, minimal and maximal total gene counts, and a minimal number of detected genes (**Table S4**). Further, genes detected in fewer than ten cells were removed. The gene expression counts were normalized per cell to 10,000 and log1p-transformed. The surface protein counts of the post-infusion sample were transformed through the centered-log-ratio (clr) across all corresponding sequencing runs.

Clonotypes were defined as cells expressing identical CDR3α and CDR3β amino acid sequences on primary or secondary chains, with clonal expansion calculated at the sample and dataset level. The donor of the post-infusion cells was annotated based on the hashtag antibody counts via HashSolo^33^ with default parameters. As the pre-infusion samples lacked hashtag antibodies, the two sequencing runs were demultiplexed separately into three singlet clusters and a doublet cluster using scSplit^34^ following the GitHub documentation. The singlet clusters’ genotypes of the pre-samples were matched to the donor-specific genotypes of the first post-infusion sequencing run based on the minimal mean absolute error of binarized gene sequence variations detected with scSplit. This annotation was validated by the overlapping clonotypes from assigned donors in pre-infusion samples to post-infusion donors determined through hashtag antibodies and coherent clonal overlap within both pre-infusion samples. Cells annotated as empty cells or doublets were removed from the dataset.

CAR^+^ cells were determined by a count greater than zero of the Hu.CD19 antibody (post-infusion only) or the CAR-gene in the transcriptome. The cells were annotated as CD8^+^, CD4^+^, CD4^+^/CD8^+^, and CD4^-^/CD8^-^ (DN) through thresholds on the cells’ gene expression level of CD8A, CD8B, and CD4 (>0.3) as well as the clr-transformed surface protein markers Hu.CD8 (>1.5) and Hu.CD4_RPA.T4 (>1). γδ-T cells were defined by gene expression of *TRGC1, TRGC2*, or *TRDC* greater than zero.

The UMAP^35^ representation at 15 neighbors and the Leiden clustering^36^ at a resolution of 2.0 were calculated on the top 5,000 highly variable genes, excluding TCR-forming genes and the CAR gene. The reactive cluster of the pre-infusion sample was determined by a high fraction of stimulated CAR^+^ cells as well as elevated *IFNG* and contained clonotypes were annotated as reactive. Differentially expressed genes and surface protein markers were determined via ‘scanpy.tl.rank_genes_groups’ using a t-test with Benjamini-Hochberg correction, using the log1p- and clr-transformed UMI counts, respectively.

### Data and Code Availability

The raw data of the single-cell sequencing experiments are publicly accessible at NCBI GEO under the accession numbers GSE299415 (pre-infusion) and GSE299416 (post-infusion). The processed data containing both samples are available at Zenodo (10.5281/zenodo.17018664). The analysis code can be accessed at: https://github.com/SchubertLab/CAR_tcell_study.

### Genetic engineering of CAR-T cells with transgenic TCRs (CAR-TCR cells)

For the generation of CAR-TCR cells, CAR transduction was combined with orthotopic TCR replacement (OTR). OTR was largely performed as described previously^20,21^. A brief description with all relevant alterations to the published protocol is provided below.

### Transgenic TCR DNA template design

The DNA template was designed *in silico* and synthesized by Twist Bioscience as a sequence-verified plasmid gene. The construct had the following structure: The left homology arm (LHA) and a self-cleaving peptide P2A was followed by the TCR β-chain comprising the human variable part (VDJβ) and the murine TCR β-constant region with an additional cysteine bridge (*mTRBC*-Cys)^37^. The subsequent self-cleaving peptide T2A separated the β-chain from the following α-chain which consisted of the human variable part (VJα) followed by the murine TCR α-constant region with an additional cysteine bridge (*mTRAC*-Cys)^37^. After the stop codon and the bovine growth hormone polyA signal, the right homology arm (RHA) concluded the homology-directed repair (HDR) template. For sequences of these segments see **Table S5**.

### Double-stranded DNA production

The lyophilized plasmid was reconstituted with PCR-grade water and amplified by PCR to generate a linearized double-stranded HDR template. A truncated Cas9 Target Sequence (tCTS) was incorporated at the 5’-end of the HDR template by the forward primer of the PCR reaction. Each 50 µL PCR reaction contained 1x Herculase II Reaction Buffer, 0.4 µM *hTRAC* HDR tCTS LHA genomic forward primer targeting the LHA (5’-TCTCTCTCTCAGCTGGTACACGGCTGCCTTTACTCTGCC AGAG-3’), 0.4 µM *hTRAC* HDR genomic reverse primer targeting the RHA (5’-CATCATTGACCAGAGCTCTG-3’), 0.5 mM dNTPs, 2 µL Herculase II Fusion DNA Polymerase, and 60 ng reconstituted DNA. The PCR was run with the following cycling conditions: Initial denaturation at 95°C for 3 min, 34 cycles of 95°C for 30 sec, 63°C for 30 sec and 72°C for 3 min, final elongation at 72°C for 3 min. Successful amplification was confirmed with a 1% agarose gel and amplified HDR template was purified with a MinElute PCR Purification Kit (Qiagen, 28004) according to the manufacturer’s instructions.

### T-cell activation for genetic editing

PBMCs were isolated from blood provided by healthy volunteers after their informed consent (Transfusion Medicine, University Hospital Erlangen) and cryopreserved at -80°C for storage. Ethics approval was granted by the local Ethics Committee of the Medical Faculty of the University Hospital of Erlangen, Friedrich-Alexander Universität Erlangen-Nürnberg, Germany (392_20Bc). For T-cell activation, PBMCs were thawed and overnight-rested at 2×10^6^ cells/mL in cRPMI medium supplemented with 50 U/mL IL-2 (Peprotech, 200-02). Afterwards, 1×10^6^ cells/mL PBMCs were activated for two days at 37°C, 5% CO_2_ on tissue-culture flasks coated with 1 µg/mL anti-CD3 (BioLegend, 317302) and 1 µg/mL anti-CD28 antibodies (BioLegend, 302902) in cRPMI medium supplemented with 300 U/mL IL-2, 5 ng/mL Interleukin-7 (Peprotech, 200-07), and 5 ng/mL Interleukin-15 (Peprotech, 200-15).

### Ribonucleoprotein (RNP) production

RNPs targeting the endogenous *hTRAC* and *hTRBC* locus were prepared. Per electroporation sample, 3.5 µL of *hTRAC* and 3 µL of *hTRBC* RNP (final concentration 20 µM) were prepared. First, 40 µM gRNAs were produced by mixing equimolar amounts of trans-activating crRNA (tracrRNA) (Integrated DNA Technologies, 1072534) with *hTRAC* crRNA (5’-AGAGTCTCTCAGCTGGTACA-3’^8^, Integrated DNA Technologies) or *hTRBC* crRNA (5’-GGAGAATGACGAGTGGACCC-3’^20^, Integrated DNA Technologies) and incubating the mixtures at 95°C for 5 min. After cool down to room temperature, 50 µg poly-*L*-glutamic acid (PGA, Sigma-Aldrich, P4761) per sample were added to *hTRAC* gRNA^38,39^ before 20 µM electroporation enhancer (Integrated DNA Technologies, 10007805) were added to both *hTRAC* and *hTRBC* gRNA. RNP production was concluded by adding equal volume of Cas9 Nuclease V3 (Integrated DNA Technologies, 1081059, diluted to 6 µM) to *hTRAC* and *hTRBC* gRNA, respectively. RNPs were incubated for 15 min at RT and subsequently stored on ice for processing at the same day. For the calculations above, the volume of PGA was not considered.

### Orthotopic TCR replacement

6h prior to electroporation, the DNA-sensing inhibitor RU.521 (InvivoGen, inh-ru52)^38^ was added to the cells at a final concentration of 4.82 nM. Activation was stopped by transferring cells into fresh cRPMI medium. For electroporation, 1×10^6^ activated cells per electroporation sample were resuspended in 20 µL P3 electroporation buffer (Lonza, V4SP-3960) and then mixed with DNA/RNP mix (0.5 µg HDR template, 3.5 µL *hTRAC* and 3 µL *hTRBC* RNPs. For the CAR-only samples, HDR template was excluded). After transfer into the 16-well Nucleocuvette^TM^ Strip (Lonza, V4SP-3960), cells were electroporated (pulse sequence EH100) in the Lonza 4D-Nucleofector^TM^. Immediately afterwards, cells were rescued in 900 µL of antibiotic-free medium (cRPMI without antibiotics) supplemented with 180 U/mL IL-2. After 15 min, 100 µL of antibiotic-free medium containing 0.5 µM HDAC class I/II Inhibitor Trichostatin A (AbMole, M1753) and 10 µM DNA-dependent protein kinase (DNA-PK) inhibitor M3814 (chemietek, CT-M3814) were added to each sample^40^. Cells were incubated for 12-18 h in a 24-well plate, before the medium was supplemented with an antibiotic mix containing gentamicin, penicillin and streptomycin to produce cRPMI medium. To remove Trichostatin A and M3814 24 h after electroporation, cells were centrifuged (480 xg, 5 min, room temperature) and resuspended in 1 mL fresh cRPMI medium supplemented with 180 U/mL IL-2. Mock samples were not electroporated and transferred to the 24-well plate with cRPMI and 180 U/mL IL-2 directly after activation.

### Transduction of CAR

To generate CAR-TCR, CAR-only or CAR-endoTCR cells, respectively, TCR-only, TCR-KO or mock cells were transduced with a CAR (JCAR17-construct) comprising an anti-CD19 (FMC63) single chain variable fragment, Strep-tag hinge domain, CD28 transmembrane domain, 4-1BB co-stimulatory domain and intracellular CD3ζ signaling domain. For sequences of these segments see **Table S3**. Transduction was performed 24h after electroporation. For virus production, RD114 cells were seeded into a 6-well plate in cDMEM (DMEM Medium (Life Technologies, 11995073) + 10% heat-inactivated FCS, 0.05 mM β-mercaptoethanol, 0.05 mg/mL gentamicin, 1.1915 g/L HEPES, 0.2 g/L L-glutamine, 100 U/mL Penicillin-Streptomycin). At 60-80% confluency, the cells were transfected with the CAR-containing plasmid using jetOPTIMUS® DNA Transfection Reagents (Sartorius, 101000051) according to the manufacturer’s instructions. Briefly, per well of the 6-well plate 2 µg CAR plasmid-DNA were diluted in 200 µl of jetOPTIMUS, vortexed and spun down. After adding 2 µL jetOptimus reagent and vortexing again, the mixture was incubated for 10 min at room temperature. After medium change to DMEM without supplements, 200 µL of the transfection mixture were added dropwise to each well of RD114 cells. The cells were incubated at 37°C with 5% CO_2_ for 6 h, before the medium was replaced with cDMEM. After 48 h, the viral supernatant was collected and stored at 4°C. One day prior to transduction, a non-tissue culture treated 24-well plate was coated with 500 µL/well RetroNectin (1 µg/mL, TaKaRa Bio Europe, T100A) at 4°C overnight. Afterwards, RetroNectin was removed and wells were blocked with 500 µL PBS with 2% BSA for 30 min at room temperature and then washed twice with PBS. For transduction, 600 µL/well of viral supernatant were centrifuged onto the plate for 2 h at 2000 xg, 32°C. Meanwhile, TCR-only, TCR-KO and mock samples were resuspended in 1 mL fresh cRPMI, respectively. After supernatant removal from the plate, 500 µL of cell suspension were added per well. Cells were centrifuged for 15 min at 480 xg, 32°C. Then, 500 µL of cRPMI with 180 U/ml IL-2 were added to each well and cells were incubated at 37°C, 5% CO_2_ for 3 days.

### Evaluating engineering efficiency

On day 3 after transduction, aliquots of the cells were washed with cold FACS buffer twice. Next, cells were incubated for 20 min in the dark on ice with the following antibodies: anti-human CD4-PE-Cy7 (1:200, BD Pharmingen, 560649), anti-human CD8a-APC (1:400), anti-hTCR-FITC (1:200, BioLegend, 306706), anti-mTRBC-APC/Fire750 (1:100, BioLegend, 109246), Streptavidin-eF450 (1:100, eBioscience, 48-4317-82), anti-EGFRt-PE (1:100, BioLegend, 352903), and ZombieAqua (1:500). After two washing steps, flow cytometric analysis was performed on a LSRFortessa (BD).

### Sorting and feeder-cell expansion

Specific cell populations were isolated by FACS. Cells were collected and washed twice in cold FACS buffer. Cells were stained as described above with the same antibodies. After washing twice, cells were resuspended in 1 mL FACS buffer. CD8^+^ CAR-TCR (CD8^+^CD4^-^Streptavidin^+^EGFRt^+^hTCR^-^mTRBC^+^), CD8^+^ CAR-only (CD8^+^CD4^-^Streptavidin^+^EGFRt^+^hTCR^-^mTRBC^-^), CD8^+^ CAR-endoTCR (CD8^+^CD4^-^ Streptavidin^+^EGFRt^+^hTCR^+^mTRBC^-^) and CD8^+^ TCR-only (CD8^+^CD4^-^Streptavidin ^-^EGFRt^-^hTCR^-^mTRBC^+^) cells were sorted into pre-coated 1.5 mL tubes containing cRPMI on a MoFlo Astrios EQ sorter (Beckman). For feeder-cell expansion, cryopreserved PBMCs (feeder cells) were thawed, rested overnight and then irradiated with 35 Gy. After 30 min rest, irradiated cells were washed with cRPMI twice and resuspended to 1×10^6^ cells/mL in cRPMI. Four irradiated feeder cells were added per sorted cell and cultured in cRPMI with 180 U/mL IL-2 and 1 μg/mL phytohaemagglutinin (PHA) (Thermo Scientific, R30852801) at 37°C with addition of 180 U/mL IL-2 every 2-3 days. On day 7, sorted cells were counted, resuspended to 1×10^6^ cells/mL and then cultured in cRPMI containing 180 U/mL IL-2, 1 μg/mL PHA and fresh irradiated feeder cells (4 per sorted cell) with addition of 180 U/mL IL-2 every 2-3 days. After 5 days, sorted cells were resuspended in fresh cRPMI containing 50 U/mL IL-2. Two days later, sorted cells were co-cultured with target cells.

### Peptide-pulsing of HLA-A2^+^ K562 or Nalm6 cells

To assess the reactivity of engineered cells against the HLA-A*02– restricted CMV epitope NLVPMVATV (WT) or its altered peptide ligands NLVPMVAAV (Mut1) and NLVPAVATV (Mut2), engineered cells were co-cultured with peptide-pulsed HLA-A*02:01^+^ K562 cells. The day before the co-culture, K562 cells were irradiated at 80 Gy, subsequently washed twice with cRPMI and resuspended at 1×10^6^ cells/mL in PBS. CellTrace-FarRed staining was used to identify K562 cells in co-cultures. 1 µM CellTrace-FarRed (0.4 mM stock, Invitrogen, C34572) was added to the cell suspension and cells were incubated under agitation in the dark at room temperature. After 20 min, five times the original staining volume of cRPMI was added and suspensions were incubated for 5 min. After centrifugation, cells were resuspended at 3×10^6^ cells/mL in fresh pre-warmed cRPMI and rested for 10 min. For peptide-pulsing, stained K562 cells were pulsed with 10 µM of respective peptides overnight at 37°C, 5% CO_2_. As a negative control, K562 cells were pulsed with DMSO.

For the peptide titration experiment, K562 cells were pulsed with respective peptides at increasing concentrations (10^-11^ to 10^-5^ M), or treated with DMSO (negative control) or 25 ng/mL PMA and 1.0 µg/mL Ionomycin (positive control). Nalm6 cells were peptide-pulsed using 10 µM of respective peptides overnight at 37°C, 5% CO_2_.

### Co-culture of engineered cells with target cells

For the *trans* conditions of the co-culture experiments with K562 and Nalm6 cells simultaneously, irradiated, stained and peptide-pulsed or control K562 cells were washed three times with cRPMI, counted and resuspended to 1×10^6^ cells/mL in cRPMI. Nalm6 cells and effector cells (CAR-TCR, CAR-only or TCR-only) were washed once, counted and resuspended to 1×10^6^ cells/mL in cRPMI. Cells were cultured at effector:target:target (E:T:T) ratios of 2:1:1 or 1:2:2, as indicated. In total, 200,000 cells per sample were cultured in 200 µL cRPMI in 96-U-bottom plates for 24 h at 37°C, 5% CO_2_.

For the *cis* settings of the co-culture experiments, peptide-pulsed Nalm6 cells were washed three times with cRPMI, counted and resuspended to 1×10^6^ cells/mL in cRPMI. Effector cells were prepared as for the *trans* setting and were co-cultured with Nalm6 cells at 2:1 or 1:2 E:T ratios, as indicated, in 200 µL in a 96-U-bottom plate (200,000 cells in total).

For the co-culture experiment with Nalm6 cells without peptide-pulsing, effector cells were co-cultured with Nalm6 cells at 1:0, 1:1, 1:5, 1:25 and 0:1 E:T ratios in 200 µL in a 96-U-bottom plate (200,000 cells in total).

For peptide titrations, peptide-pulsed K562 cells and effector TCR-only cells were prepared as described above and were co-cultured at 1:2 E:T ratios in 200 µL in a 96-U-bottom plate (200,000 cells in total).

Co-cultures were analyzed by flow cytometry. 10 µL Counting Beads (Invitrogen, 01-1234-42) were added per well. Then, cells were transferred to 96-V-bottom plates and washed twice with cold FACS buffer. Cells were stained with anti-human CD19-PE/CFS594 (1:200, BD Horizon, 562294), anti-human CD8a-BUV395 (1:200, BD Horizon, 563795), Streptavidin-eF450 (1:100), anti-EGFRt-PE (1:100), anti-hTCR-FITC (1:100), anti-human CD69-PerCP/Cy5.5 (1:200, BD Pharmingen, 560738), anti-human CD137-PE-Cy7 (1:50, BioLegend, 309817), ZombieAqua (1:500) and anti-mTRBC-BV711 (1:200, BioLegend, 109243) or anti-mTRBC-APC/Fire750 (1:100, BioLegend, 109246) on ice in the dark for 20 min. Then, cells were washed twice with cold FACS buffer and acquired on a LSRFortessa (BD).

### Conjugation assay

Indicated Effector cells were resuspended to 1×10^6^ cells/mL in PBS and stained with 0.5 µM CellTrace CFSE (1.8 mM stock, Invitrogen, C34554) under agitation in the dark at room temperature. After 20 min, five times the original staining volume of cRPMI was added and incubated for 5 min. After centrifugation, cells were resuspended at 1×10^6^ cells/mL in fresh pre-warmed cRPMI and rested for 10 min. Nalm6 cells were washed with FACS buffer and stained with anti-HLA/A2-PE-Cy7 (1:100, Biolegend, 343314) for 20 min at room temperature in the dark. After washing with FACS buffer, stained Nalm6 cells were resuspended to 1×10^6^ cells/mL in pre-warmed cRPMI. K562 cells were CellTrace-FarRed-stained and peptide-pulsed as described. Labeled effector and target cells were then co-cultured *in cis* or *in trans*, as described above, at an E:T(:T) ratio of 1:2(:2) at 37°C, 5% CO_2._ After 10 min or 30 min, cells were fixed by adding 0.33% Paraformaldehyde (16% stock, Science service, E15710). Immediately after, samples were placed on ice and with as little disturbance as possible, acquired on a LSRFortessa (BD). Frequencies of effector:target doublets (effector cell and K562 or Nalm6) or triplets (effector cell and K562 and Nalm6) were analyzed with FlowJo 10.

### Confocal microscopy

K562 cells were pulsed with indicated peptides as described above, but without CellTrace-FarRed staining, and resuspended to 1×10^6^ cells/mL in cRPMI. On the day of the experiment, Nalm6 cells were stained with Cell Proliferation Dye eFluor 450 (CPD, eBioscience, 65-0842-85), to be distinguishable in the co-culture setting. Nalm6 cells were washed with PBS and resuspended in PBS at a density of 5×10^6^ cells/mL. An equal volume of PBS with 30 µM CPD was added dropwise to the cells, while slowly vortexing and cells were incubated at 37°C, 5% CO_2_ in the dark for 20 min. Five times the volume of ice-cold cRPMI was added and cells were incubated for 5 min on ice. Cells were subsequently washed 3 times with cold cRPMI, resuspended at 1×10^6^ cells/mL in cRPMI and incubated at 37°C, 5% CO_2_ for 30 min. CAR-TCR cells were resuspended at 1×10^6^ cells/mL in cRPMI. CAR-TCR cells were co-incubated with target cells at an E:T:T ratio of 2:1:1 in 500 µL cRPMI (500,000 cells in total) on 12mm-coverslips (Marienfeld, MARI0112520) coated with Poly-D-Lysine (0.05 mg per coverslip, Invitrogen, A3890401) in 24-well plates at 37°C, 5% CO_2_. To stain surface receptors, the following antibodies were added directly at the beginning of the co-culture: anti-murine TCR antibody (1:100, BioLegend, 109201), Strep-Tactin DY-488 (1:200, IBA, 2-1562-050). After 30 min, co-cultures were spun onto the plates for 3 min at 400 xg, room temperature, medium was removed, cells were washed 3 times with PBS and fixed with 4% Paraformaldehyde for 10 min at room temperature in the dark. All following steps were performed at room temperature in the dark. Fixed cells were washed twice with PBS, permeabilized with Triton-×100 (0.1%, VWR, SIALT9284-100ML) for 15 min, washed twice again and then blocked in PBS with 5% normal goat serum (Thermo Fisher, 10000C) for 1 h. Subsequently, cells were stained first with anti-perforin (1:500, Merck, ZMS1188-25UL) and then with phalloidin-AF568 or -AF647 (1:200, Thermo Fisher, A12380 or A22287), goat anti-mouse IgG AF568 or AF647 (1:500, Thermo Fisher, A-11031 or A-21235) and goat anti-armenian hamster IgG AF750 (1:100, abcam, ab175760) in 5% normal goat serum in PBS for 1 hour. After each antibody incubation, cells were washed 3 times with 5% normal goat serum in PBS for 5 min each. Control co-cultures only stained for one marker were prepared as staining controls. Coverslips were mounted onto glass slides (VWR, 630-1985) in ProLong Gold Antifade Mountant (Fisher Scientific, P10144). Images were acquired on a Leica Stellaris 8 in Lightning mode using a 63x/1.3 NA glycerol objective. Images of CAR-TCR cells interacting with Nalm6 and K562 cells only or with both target cells simultaneously were acquired from the same co-culture setting as 3D stacks. Regions of interest were chosen based on occurrence of effector:target:target triplets and all triplets that we found were acquired. Triplets were defined based on direct physical contact between a big cell only showing phalloidin signals (K562), a smaller CPD-labeled cell (Nalm6) and a smaller cell with TCR and CAR signals (effector). Perforin signals were not considered for region of interest choice. Acquired images were automatically deconvolved in the Leica Application Suite X.

### Confocal image analysis

Confocal images shown here were deconvolved with Huygens (Scientific Volume Imaging) and prepared with Fiji^41^. Colocalization analyses (Pearson’s correlation coefficients and Manders’ coefficients) were performed on automatically deconvolved images from the microscope software with the JACoP v2.0^42^ plugin on regions of interest covering whole individual effector cells in the indicated effector:target interactions. Thresholds for specific signals for the analysis were determined based on single staining samples for all used antibodies and reagents. Costes randomization was used to validate non-random colocalization and reached Costes p-values of >95% for all analyses. To quantify signal intensities (mean gray values) in the immunological synapses, 3D TCR and CAR synapse regions of the effector cells were cropped based on receptor and actin signal density and mean fluorescence intensities within the cropped images were measured with the 3D objects counter (intensity threshold 0). As control, signal intensities in non-synaptic regions were measured. Signal intensities were normalized by subtracting signals within the non-synaptic region from signals within synaptic regions, where indicated. Signal intensities in the line graph in Fig. 3C were measured with the ‘Plot Profile’ function of Fiji in the indicated region of the sum intensity projection in Fig. 3B. Images represent sum intensity projections across 3D z stacks, unless indicated otherwise, and the lookup tables ‘Magenta Hot’ (TCR), ‘Green Fire Blue’ (CAR) and ‘Yellow Hot’ (perforin) were applied, where indicated, to better visualize differences in signal intensities.

### Data analysis and visualization

Data graphs were generated with GraphPad Prism 10. For statistical testing, data were first tested for normal distribution, then the appropriate test was chosen and is highlighted in the figure legends. In all graphs, only statistically significant results are highlighted. In violin plots, solid lines indicate median, dashed lines indicate 25^th^ and 75^th^ percentiles. Schemes and figures were generated with Affinity Designer (Serif (Europe) Ltd, version 1.10.1).

## Author contributions

Conceptualization: K.S, data curation: C.M, F.D., E.S.; formal analysis: C.M, F.D., E.S.; funding acquisition: K.S., B.S.; investigation: C.M, F.D., E.S., A.P., M.B., V.R., M.S.; methodology: C.M., M.B., E.S.; project administration: K.S., resources: S.V., D.M., A.M, S.U., D.H.B., H.B., K.S.; software: F.D., B.S.; supervision: K.S., N.P.K., M.H., B.S., D.M., A.M; validation: C.M., F.D., E.S.; visualization: C.M., F.D., E.S., K.S.; writing original draft: K.S., C.M., F.D., E.S., N.P.K.; review & editing: all authors.

## Acknowledgments

This work was mainly supported by grants from the German Federal Ministry for Research, Technology and Space (BMFTR, projects 01KI2013, 01KI2516 and 031L0290B) to K.S.; the BMFTR (project 031L0290A) to B.S. Further support came from grants by the Else Kröner-Fresenius-Stiftung (project 2020_EKEA.127) and the German Research Foundation (DFG) through the research training group RTG 2504 (project 401821119), the Heisenberg program (project SCHO 1949/1-1), the TRR221/3 (324392634, project A09) and the SFB 1755 CASCAID (550296805; project #13), the Aventis Foundation (project 2024: Life Science Bridge Award), as well as the Yellow4FLAVI project, funded by the European Union, under the Horizon Europe Programme (Grant Agreement N°101137459) to K.S. F.D. is supported by Helmholtz Munich’s Health AI Fellowship and acknowledges financial support from the Joachim Herz Stiftung. Funding agencies had no influence on the study design or implementation. The study was funded by the European Union. Views and opinions expressed are, however, those of the authors only and do not necessarily reflect those of the European Union or HADEA.

We thank all study participants for making this work possible. We thank members from the Schober laboratory for experimental support and critical discussion. We thank Johannes Huppa (Berlin) for feedback on the manuscript, thank Dimitrios Wagner (Berlin/Houston) and Jonas Kath (Berlin) for advice on T-cell engineering optimization and Dirk Mielenz (Erlangen) for advice on immune synapse imaging. Furthermore, we gratefully acknowledge the generous support by the Manfred Roth-Stiftung, Fürth, Germany, and thank the Core Unit Cell Sorting and Immunomonitoring Erlangen and the Optical Imaging Center Erlangen.

## Supplementary Materials

Figs. S1 to S3; Tables S1 to S3

## References

1. Schett, G., Mackensen, A. & Mougiakakos, D. CAR T-cell therapy in autoimmune diseases. Lancet 6736, 1–11 (2023).

2. Guedan, S., Ruella, M. & June, C. H. Emerging Cellular Therapies for Cancer. Annu. Rev. Immunol. 37, 145–171 (2018).

3. Sheih, A. et al. Clonal kinetics and single-cell transcriptional profiling of CAR-T cells in patients undergoing CD19 CAR-T immunotherapy. Nat. Commun. 11, 219 (2020).

4. Wilson, T. L. et al. Common Trajectories of Highly Effective CD19-Specific CAR T Cells Identified by Endogenous T-cell Receptor Lineages. Cancer Discov. 12, 2098–2119 (2022).

5. Louie, R. H. Y. et al. CAR+ and CAR-T cells share a differentiation trajectory into an NK-like subset after CD19 CAR T cell infusion in patients with B cell malignancies. Nat. Commun. 14, (2023).

6. Guerrero-Murillo, M. et al. Integrative single-cell multi-omics of CD19-CARpos and CARneg T cells suggest drivers of immunotherapy response in B cell neoplasias. Cell Reports Med. 5, 101803 (2024).

7. Stenger, D. et al. Endogenous TCR promotes in vivo persistence of CD19-CAR-T cells compared to a CRISPR/Cas9-mediated TCR knockout CAR. Blood 136, 1407–1418 (2020).

8. Ren, J. et al. Multiplex Genome Editing to Generate Universal CAR T Cells Resistant to PD1 Inhibition. Clin. Cancer Res. 23, 2255–2266 (2017).

9. Ghosh, A. et al. Donor CD19 CAR T cells exert potent graft-versus-lymphoma activity with diminished graft-versus-host activity. Nat. Med. 23, 242–249 (2017).

10. Li, C.-H. et al. Long-term outcomes of GD2-directed CAR-T cell therapy in patients with neuroblastoma. Nat. Med. (2025) doi:10.1038/s41591-025-03513-0.

11. Cruz, C. R. Y. et al. Infusion of donor-derived CD19-redirected virus-specific T cells for B-cell malignancies relapsed after allogeneic stem cell transplant: a phase 1 study. Blood 122, 2965–2973 (2013).

12. Pule, M. A. et al. Virus-specific T cells engineered to coexpress tumor-specific receptors: Persistence and antitumor activity in individuals with neuroblastoma. Nat. Med. 14, 1264–1270 (2008).

13. Kondo, T. et al. Engineering TCR-controlled fuzzy logic into CAR T cells enhances therapeutic specificity. Cell 188, 2372-2389.e35 (2025).

14. Uslu, U., Schuler, G., Dörrie, J. & Schaft, N. Combining a chimeric antigen receptor and a conventional T-cell receptor to generate T cells expressing two additional receptors (TETARs) for a multi-hit immunotherapy of melanoma. Exp. Dermatol. 25, 872–879 (2016).

15. Teppert, K. et al. CAR’TCR-T cells co-expressing CD33-CAR and dNPM1-TCR as superior dual-targeting approach for AML treatment. Mol. Ther. Oncol. 32, 200797 (2024).

16. Wachsmann, T. L. A. et al. CAR-mediated target recognition limits TCR-mediated target recognition of TCR- and CAR-dual-receptor-edited T cells. Mol. Ther. 33, 1642– 1658 (2025).

17. Yang, Y. et al. TCR engagement negatively affects CD8 but not CD4 CAR T cell expansion and leukemic clearance. Sci. Transl. Med. 9, 1–13 (2017).

18. Fischer, D. S. et al. Single-cell RNA sequencing reveals ex vivo signatures of SARS-CoV-2-reactive T cells through ‘reverse phenotyping’. Nat. Commun. 12, 4515 (2021).

19. Melenhorst, J. J. et al. Decade-long leukaemia remissions with persistence of CD4+ CAR T cells. Nature 602, 503– 509 (2022).

20. Schober, K. et al. Orthotopic replacement of T-cell receptor α- and β-chains with preservation of near-physiological T-cell function. Nat. Biomed. Eng. 3, 974–984 (2019).

21. Moosmann, C., Müller, T. R., Busch, D. H. & Schober, K. Orthotopic T-cell receptor replacement in primary human T cells using CRISPR-Cas9-mediated homology-directed repair. STAR Protoc. 3, 101031 (2022).

22. Barden, M. et al. CAR and TCR form individual signaling synapses and do not cross-activate, however, can co-operate in T cell activation. Front. Immunol. 14, 1–13 (2023).

23. Drost, F. et al. Predicting T cell receptor functionality against mutant epitopes. Cell Genomics 4, 100634 (2024).

24. Gattinoni, L., Speiser, D. E., Lichterfeld, M. & Bonini, C. T memory stem cells in health and disease. Nat. Med. 23, 18–27 (2017).

25. Sommermeyer, D. et al. Chimeric antigen receptor-modified T cells derived from defined CD8+ and CD4+ subsets confer superior antitumor reactivity in vivo. Leukemia 492– 500 (2015) doi:10.1038/leu.2015.247.

26. Fraietta, J. A. et al. Determinants of response and resistance to CD19 chimeric antigen receptor (CAR) T cell therapy of chronic lymphocytic leukemia. Nat. Med. 24, 563–571 (2018).

27. Chen, G. M. et al. Integrative Bulk and Single-Cell Profiling of Premanufacture T-cell Populations Reveals Factors Mediating Long-Term Persistence of CAR T-cell Therapy. Cancer Discov. 11, 2186–2199 (2021).

28. Wiedemann, A., Depoil, D., Faroudi, M. & Valitutti, S. Cytotoxic T lymphocytes kill multiple targets simultaneously via spatiotemporal uncoupling of lytic and stimulatory synapses. Proc. Natl. Acad. Sci. U. S. A. 103, 10985–90 (2006).

29. McGavern, D. B., Christen, U. & Oldstone, M. B. A. Molecular anatomy of antigen-specific CD8(+) T cell engagement and synapse formation in vivo. Nat. Immunol. 3, 918–25 (2002).

30. Heumos, L. et al. Best practices for single-cell analysis across modalities. Nat. Rev. Genet. 24, 550–572 (2023).

31. Wolf, F. A., Angerer, P. & Theis, F. J. SCANPY: large-scale single-cell gene expression data analysis. Genome Biol. 19, 15 (2018).

32. Sturm, G. et al. Scirpy: a Scanpy extension for analyzing single-cell T-cell receptor-sequencing data. Bioinformatics 36, 4817–4818 (2020).

33. Bernstein, N. J. et al. Solo: Doublet Identification in Single-Cell RNA-Seq via Semi-Supervised Deep Learning. Cell Syst. 11, 95-101.e5 (2020).

34. Xu, J. et al. Genotype-free demultiplexing of pooled single-cell RNA-seq. Genome Biol. 20, 290 (2019).

35. McInnes, L., Healy, J., Saul, N. & Großberger, L. UMAP: Uniform Manifold Approximation and Projection. J. Open Source Softw. 3, 861 (2018).

36. Traag, V. A., Waltman, L. & van Eck, N. J. From Louvain to Leiden: guaranteeing well-connected communities. Sci. Rep. 9, 5233 (2019).

37. Cohen, C. J. et al. Enhanced antitumor activity of T cells engineered to express T-cell receptors with a second disulfide bond. Cancer Res. 67, 3898–3903 (2007).

38. Kath, J. et al. Pharmacological interventions enhance virus-free generation of TRAC-replaced CAR T cells. Mol. Ther. Methods Clin. Dev. 25, 311–330 (2022).

39. Nguyen, D. N. et al. Polymer-stabilized Cas9 nanoparticles and modified repair templates increase genome editing efficiency. Nat. Biotechnol. 38, 44–49 (2020).

40. Shy, B. R. et al. High-yield genome engineering in primary cells using a hybrid ssDNA repair template and small-molecule cocktails. Nat. Biotechnol. 41, 521–531 (2023).

41. Schindelin, J. et al. Fiji: an open-source platform for biological-image analysis. Nat. Methods 9, 676–82 (2012).

42. Bolte, S. & Cordelières, F. P. A guided tour into subcellular colocalization analysis in light microscopy. J. Microsc. 224, 213–32 (2006).

