## Supplementary Figures for "Antigen receptor competition in CAR-TCR cells changes cytotoxicity outcomes upon spatial antigen separation"

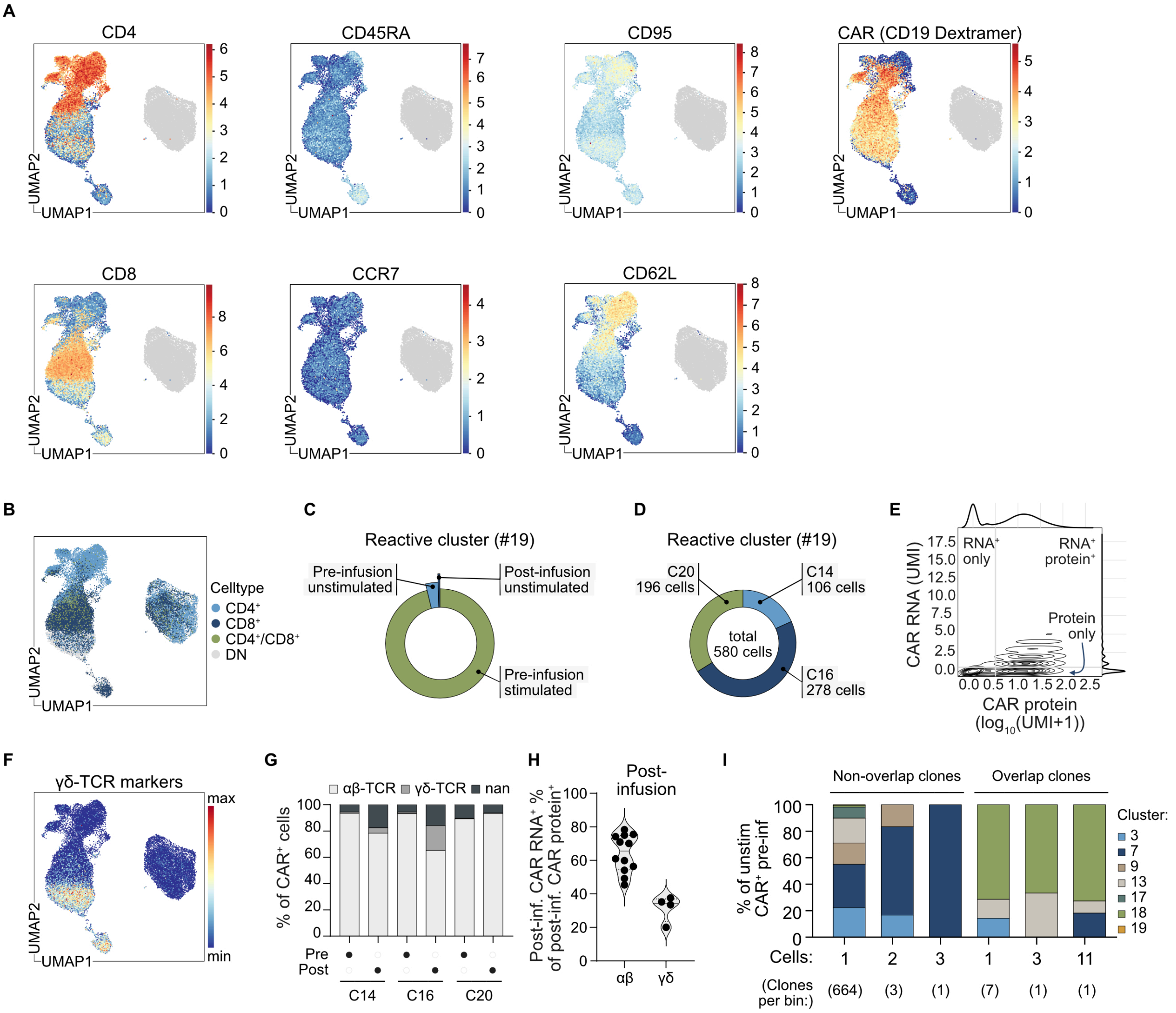


**Figure S1: Additional single-cell analysis of pre- and post-infusion CAR-T cells and clonotypes. (A)** Surface marker expression in post-infusion sample as detected by CITE-seq antibodies (anti-CD4, anti-CD8, anti-CD45RA, anti-CCR7, anti-CD95 and anti-CD62L) and CD19-dCODE-Dextramers. Expression levels are shown as ln(UMI+1). **(B)** UMAP showing $CD4^+$, $CD8^+$, double positives ($CD4^+/CD8^+$) and double negatives (DN) cells. **(C-D)** Stimulation condition (C) and donor distribution (D) of cells within the reactive cluster 19. **(E)** Single-cell expression levels of CAR protein ($\log_{10}$(UMI+1) and CAR RNA (UMI) of post-infusion cells. **(F)** Expression of γδ-TCR markers *TRDC*, *TRGC2* and *TRGC1*. **(G)** Distribution of $CAR^+$ α/β-TCR cells, γδ-TCR cells and rest (“nan”) pre- and post-infusion per donor. **(H)** CAR $RNA^+$ among CAR $protein^+$ cells in post-infusion clusters. For “α/β-TCR” clusters (clusters 1, 2, 5, 6, 8, 12, 14, 15, 16, 20, 21, 22) only α/β-TCR cells and for “γδ-TCR” clusters (clusters 0, 4, 10, 11) only γδ-TCR cells as defined in (G) were considered. Each dot represents one Leiden cluster. **(I)** Bin analysis of Leiden cluster distribution of unstimulated $CAR^+$ pre-infusion cells of non-overlapping and overlapping $CAR^+$ clones. Clone sizes and number of clones belonging to the respective bins are indicated.

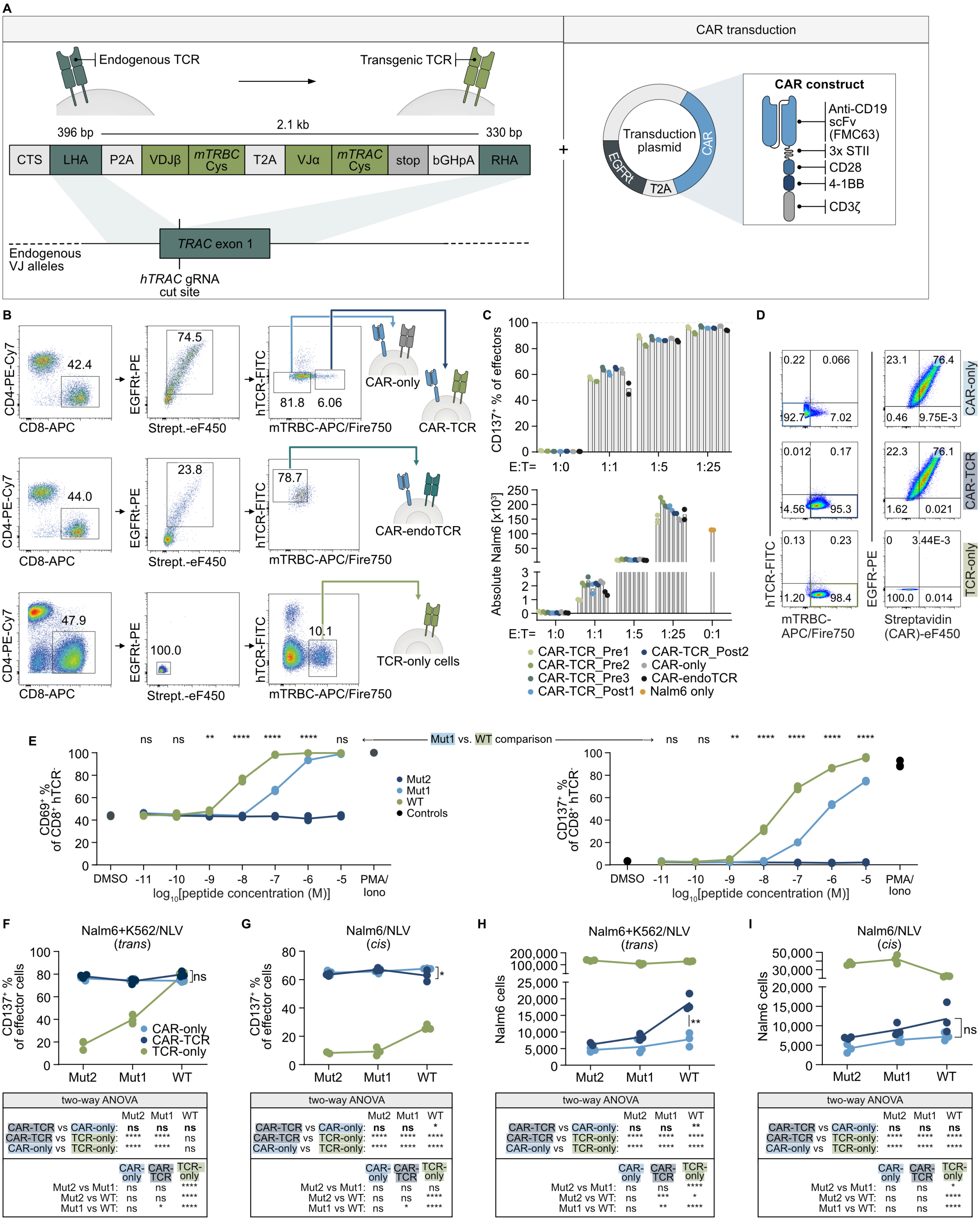


**Figure S2: Application of the CAR-TCR platform to investigate the influence of TCR presence and clonotype on CAR-T cell functionality. (A)** Genetic information used for the CAR-TCR platform combining orthotopic TCR replacement (OTR) and CAR transduction. DNA template for TCR re-expression included transgenic TCR α-and β-chain and allows integration into the human *TRAC* locus via homology-directed repair. The transduction plasmid comprises a modified second-generation CAR construct (an anti-CD19 single-chain variable fragment (scFv), a Strep-tag hinge-domain, a CD28-derived transmembrane domain, a 4-1BB costimulatory domain and a CD3ζ signaling domain) followed by a T2A element and a truncated epidermal growth factor receptor (EGFRt, used to identify transduced cells). **(B)** Gating strategy for sorting of edited effector cells. Cells were pre-gated on single, live lymphocytes. Data for CAR-only, CAR-TCR and CAR-endoTCR cells are representative of two independent experiments and data for TCR-only cells are representative of two different, independent experiments. **(C)** Percentage of CD137$^+$ effector cells (top) and absolute Nalm6 target cells (bottom) following 24 h co-culture of Nalm6 cells with indicated effector cells at different effector:target (E:T) ratios. TCRs were selected from the pre-infusion product (top 3 most expanded exclusively CAR$^+$ TCR clonotypes, strictly not re-appearing in the post-infusion sample, for which not a single cell was recruited into the reactive cluster: Pre1, Pre2 and Pre3) or the post-infusion product (top 2 most expanded, purely CAR$^+$ clonotypes: Post1, Post2). **(D)** Purity of effector cells after sorting. Pregated on live, single $CD8^+$ lymphocytes (left) or the respective parent population from left (right). **(E)** Upregulation of activation markers CD69 (left) and CD137 (right) in TCR-only cells following 24 h co-culture with K562 cells treated with DMSO (negative control), pulsed with increasing amounts of Mut2, Mut1 or WT peptide or treated with PMA/Ionomycin (positive control). **(F-I)** Percentage of $CD137^+$ effector cells (F,G) and absolute numbers of Nalm6 cells (H,I) following 24 h of co-culture of effector cells with peptide-pulsed K562 cells and Nalm6 cells (*trans*, F,H) or peptide-pulsed Nalm6 (*cis,* G,I) at an E:T(:T) of 1:2(:2). Bars (C) and lines (E-I) show means. Data points represent technical replicates (n=2 (C), n=3 (E-I)) of one experiment. Statistical testing: two-way ANOVA followed by Šídák's multiple comparisons test (E) or Tukey's multiple comparisons test (F-I). Significance: * $p<0.05$, ** $p<0.01$, *** $p<0.001$, **** $p<0.0001$.

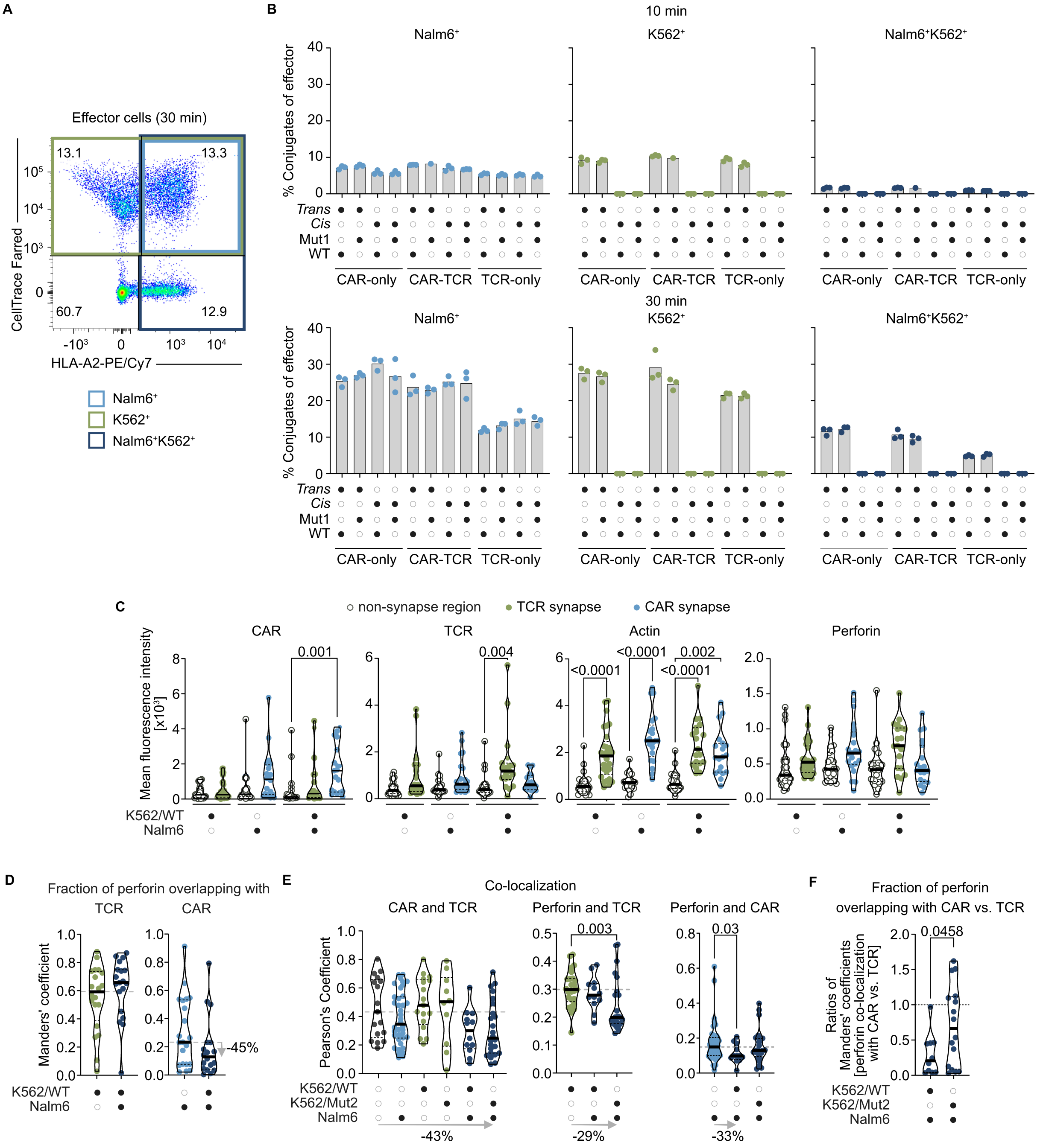

**Figure S3: Additional analysis of CAR- and TCR-interference in CAR-TCR cells on cellular interactions and immunological synapse formation. (A-B)** Representative gating (A) and quantification (B) for flow cytometric analysis of effector cells forming conjugates with K562 and/or Nalm6 cells after 10 min (B, top) or 30 min (B, bottom) of *cis* (E:T=1:2) or *trans* (E:T:T=1:2:2) co-cultures with Mut1- or WT-pulsed target cells. **(C)** Signal intensity of components in the immunological synapses of CAR-TCR cells interacting with indicated target cells after 30 min of *trans* co-culture with Nalm6 and WT NLV peptide-pulsed K562 cells analyzed from confocal microscopy images. Signals were measured within isolated synapse or non-synaptic regions. Statistical test: Kruskal-Wallis test followed by Dunn's multiple comparisons test. **(D-F)** Co-localization analysis of indicated synapse components from confocal microscopy images of whole CAR-TCR cells interacting with indicated target cells. K562 cells were either pulsed with WT or Mut2 NLV peptide as indicated. (D-E) Percentages at the bottom or right are calculated as the signal loss between medians of effector:Nalm6:K562 triplets as indicated and the reference indicated by the start of the arrow. (F) Ratios of Manders' coefficients quantify perforin co-localization with CAR relative to TCR. Statistical test: Mann-Whitney test (E), unpaired t test (F). Data points represent technical replicates (n=1 for CAR-TCR *trans* Mut1 10 min or n=3 for all others in B) or individual CAR-TCR cells (n=29 (C, effector:K562), n=21 (C-D, effector:Nalm6), n=19 (C-D, effector:Nalm6:K562), n=20 (D, effector:K562), n=17 (E, effector-only), n=38 (E, effector:Nalm6), n=17 (E, effector:K562/WT), n=10 (E-F, effector:K562/Mut2), n=12 (E, effector:Nalm6:K562/WT), n=24 (E, effector:Nalm6:K562/Mut2), n=16 (F, effector:Nalm6:K562/Mut2)) of one experiment representative of n=2 (B-D) or n=1 (E-F) experiments.
